# Oceanic salt spray and herbivore pressure contribute to local adaptation of coastal perennial and inland annual ecotypes of the Seep Monkeyflower (*Mimulus guttatus*)

**DOI:** 10.1101/523902

**Authors:** Damian Popovic, David B. Lowry

## Abstract

Identifying the environmental factors responsible for natural selection across different habitats is crucial for understanding the process of local adaptation. Despite its importance, only a few studies have successfully isolated the environmental factors driving local adaptation in nature. In this study, we evaluated the agents of selection responsible local adaptation of the monkeyflower *Mimulus guttatus* to coastal and inland habitats in California. We implemented a manipulative field reciprocal transplant experiment at coastal and inland sites, where we excluded aboveground stressors in an effort to elucidate their role in the evolution of local adaptation. We found that excluding these stressors, most likely a combination of salt spray and herbivory, completely rescued inland plant fitness when transplanted to coastal habitat. In contrast, the exclosures in inland habitat provided limited fitness benefit for either coastal or inland plants. We have previously established that low soil water availability belowground is the most important agent of selection in inland habitat. Therefore, our study demonstrates that a distinct set of selective agents are responsible for local adaptation at opposite ends of an environmental gradient.

## INTRODUCTION

The living world is made rich with varied biological diversity. Much of that diversity is the result of natural selection acting upon variation in wild populations. Local adaptation to differing habitats plays a key role in the evolution of morphologically, physiologically, and phenologically distinct intraspecific populations (Clausen 1951; Schemske 2000; Coyne and Orr 2004). Local adaptation can be described as divergence due to contrasting environmental conditions across a species range. This often results in a tradeoff, where home populations have higher fitness than foreign populations in each habitat (Van Tienderen 1997; Kawecki and Ebert 2004). Over time, local adaptation can lead to the evolution of prezygotic and postzygotic reproductive isolating barriers among populations (Rundle 2002; Nosil 2007; Sobel *et al*. 2010). One of the most effective barriers inhibiting introgression among plants is that of ecogeographic isolation, described as the allopatric distribution of populations enforced by local adaptation to divergent ecological and edaphic regimes (Schemske 2000; Ramsey *et al*. 2003; Husband and Sabara 2004; Kay 2006; Sobel 2014). Strong divergent selection and low relative gene flow across a species range can result in the evolution of disparate ecotypes, groups of locally adapted populations that exhibit reproductive isolation, but not to the point that they would be considered separate biological species (Lowry 2012). While local adaptation is now viewed as a cornerstone of the evolution of biological diversity, surprisingly few studies have identified the key environmental variables, or selective agents, that drive its evolution.

Almost a century has passed since Göte Turesson first introduced the concept of ecotypes (Turesson 1922). Since then, various landmark studies have introduced and popularized the use of reciprocal transplant common gardens to determine the prevalence of ecotypes in numerous species (reviewed in Leimu & Fischer 2008; Hereford 2009). Most of these inquiries have primarily focused on testing the hypothesis that the ecotypes are local adapted, while little work has been done to identify the causative selective agents (Wadgymar *et al*. 2017). Typically, studies of local adaptation have made use of field or laboratory findings and an understanding of regional natural history to make predictions about potential selective agents, whether they be herbivore resistance in aspect-specific stands of *Quercus rubra* (Sork *et al*. 1993), winter temperatures in natural populations of *Arabidopsis thaliana* (Ågren and Schemske 2012), or predator avoidance via substrate crypsis in *Chaetodipus intermedius* (Hoekstra *et al*. 2005). However, many of these works have appropriately expressed caution in inferring selective agents without employing direct experimental manipulation in field reciprocal transplants. Wadgymar *et al*. (2017) recently surveyed the local adaptation literature for studies that identified agents of local adaptation in nature through manipulative field experiments and identified just four such studies (Williamson *et al*. 1997; Bischoff *et al*. 2006; Liancourt *et al*. 2013; Maes *et al*. 2014). Only with further manipulative field experiments can we begin to elucidate the broader causal associations between environmental selective agents and local adaptation (Cheplick 2015).

In this study, we conducted a manipulative field experiment to better understand the environmental variables contributing to local adaptation in the Seep Monkeyflower, *Mimulus guttatus*. Native to Western North American, *M. guttatus* has proven valuable to the study of ecological and evolutionary genetics (Wu *et al*. 2008; Lowry and Willis 2010; Friedman *et al*. 2014; Ferris *et al*. 2016; Gould *et al*. 2017; Troth *et al*. 2018). Local adaptation has previously been demonstrated among disparate ecotypes of *M. guttatus* across California’s coast-inland moisture gradient (Hall and Willis 2006; Lowry *et al*. 2008; Lowry & Willis 2010). Inland populations endemic to ephemeral streams and seeps exhibit an annual life history, prioritizing seed production as a strategy to escape the seasonal summer drought (Vickery 1952; Lowry *et al*. 2008). In contrast, coastal populations have adopted a perennial life history, persisting year-round in long-lived headland seeps under cooler maritime conditions (Vickery 1952). Due to delayed reproductive maturity, plants of the coastal ecotype transplanted to inland habitats fail to flower before the onset of the hot summer drought (Lowry *et al*. 2008; 2010). Though avoidance of the seasonal summer drought is likely the most important factor responsible for homesite advantage of inland populations, the agents of selection underlying homesite advantage in coastal perennial populations have not been explicitly tested.

Recent work by Kooyers *et al*. (2017) revealed a potent tradeoff between growth rate and phenylproponoid glycoside (PPG) production (a vital class of herbivore resistance phytochemicals) in variable populations of *M. guttatus* across an altitudinal gradient. Paired with evidence that coastal perennial populations generally produce higher relative concentrations of PPGs that reduce herbivory (Holeski *et al*. 2013; Rotter et al. 2018), this could implicate the role of differential herbivore pressure as a biotic agent influencing divergent selection. Additionally, coastal populations have also been found to be more tolerant to salt spray (Lowry *et al*. 2008, 2009), a ubiquitous abiotic stressor in coastal habitats. Despite higher levels of herbivore resistance and salt tolerance of coastal perennial populations of *M. guttatus*, it is unknown how much impact those variables have on fitness in the nature.

Here, we conducted a manipulative reciprocal transplant experiment to investigate whether the manipulations of a habitat’s aboveground selective agents can restore fitness in maladapted foreign ecotypes. Exclosures can buffer plants from the detrimental effects of herbivory, salt spray, wind, and adverse temperatures while holding edaphic soil characteristics constant. To narrow the list of candidate selective agents affecting fitness in the field, plots at both coastal and inland sites were protected with agrofabric exclosures. We then evaluated the fitness of both ecotypes at the end of the growing season, comparing relative performance of exclosure replicates to their controlled counterparts. This design allowed us to demonstrate that edaphic soil factors are not among those agents selecting against inland *M. guttatus* recruits in coastal habitat. Rather, some combination of aboveground agents, most likely vegetative herbivory and salt exposure, are crucial for local adaptation of the coastal populations to their native habitat.

## MATERIALS AND METHODS

### Genotype selection and growth conditions

To determine what combination of selective agents contributes to differential performance in divergent *M. guttatus* genotypes along California’s coast-inland moisture gradient, we conducted a manipulative reciprocal transplant experiment. To verify that these effects are replicable within ecotypes, we used accessions from two coastal perennial (SWB and MRR) and two inland annual populations (LMC and OCC). All seeds were derived from a single field mother per accession. LMC and SWB were collected in Mendocino County, CA and have been shown to be locally adapted through previous reciprocal transplant experiments (Lowry *et al*. 2008; Lowry & Willis 2010). OCC and MRR were collected from Sonoma County, CA and have not been previously utilized in field reciprocal transplant experiments (Table 1). Seeds from each accession were gathered from the wild in previous years and stored in the Lowry Lab at Michigan State University (MSU). All accessions were grown at least one generation at the MSU greenhouse facilities to control for potential maternal effects and bulk seed stores. Since seed bulking can result in multiple generations of inbreeding in cataloged accessions, we chose among those inbred fewer than four generations (Table 1). Though some inbreeding depression was unavoidable, this screening allowed us to confidently negate the worst effects from our study.

**Table 1.** Geographic locations and inbreeding information of populations used in this study.

| Ecotype | Pop ID | Inbred | Location | Latitude (N) | Longitude (W) |
| --- | --- | --- | --- | --- | --- |
| coastal | SWB-11-1 | 1 Gen | Irish Beach, Mendocino Co., CA | 39° 02' 09" | 123° 41' 25" |
|  | MRR-13-2 | 1 Gen | Jenner, Sonoma Co., CA | 38° 27' 38" | 123° 08' 45" |
| inland | LMC-24 | 3 Gens | Yorkville, Mendocino Co., CA | 38° 51' 50" | 123° 05' 02" |
|  | OCC-31 | 2 Gens | Occidental, Sonoma Co., CA | 38° 24' 57" | 122° 56' 13" |

Seeds were sown at UC Berkeley’s greenhouse facilities on February 1^st^ 2017. Each accession was sown as a lawn upon corresponding potting flats (54.28 cm L x 27.94 cm W x 6.20 cm H) filled with Sun Gro Horticulture’s Sunshine Mix #1 (two trays per accession, eight in total), moistened prior to sowing with deionized (DI) water. Several hundred seeds were sown to ensure that enough germinated for the experiment. Each of the resulting eight flats were subsequently misted with DI water and stored in a cold room at 4°C to stratify. Coastal flats were relocated to the greenhouse after 10 days of stratification, while their inland counterparts remained for an additional week (17 days). Considering the rapid pace at which inland annuals mature relative to coastal perennials, staggering their relocation allowed us to align the life stages in all our genotypes – regardless of ecotype – for planting in the field. Seedlings were germinated under constant conditions, misted daily, and exposed to 16 hours of daylight. All flats were transported from UC Berkeley’s greenhouse facilities to the greenhouses at UC Davis’ Bodega Marine Laboratory & Reserve on February 28^th^ 2017.

### Reciprocal transplant design

To test whether site-specific agents of natural selection select against non-native ecotypes, a reciprocal transplant common garden experiment was planted along California’s coast-inland moisture gradient. Ideal coastal and inland sites were selected at two ecological reserves in Sonoma County, CA. Both gardens were planted in seeps inhabited by native populations of *M. guttatus*. The coastal garden was planted along a perennial seep at the southern end of Horseshoe Cove (Latitude: 38.315716°, Longitude: −123.068625°; 60.75 m from the ocean) on land managed by the UC Davis Bodega Marine Reserve (BMR) in Bodega Bay, CA (Figure 1). Our inland site was located along the margins of an ephemeral hillside seep (Latitude: 38.575545°, Longitude: −122.700851°; 39.84 km from the ocean) at the Pepperwood Preserve near Santa Rosa, CA. We established three split-plots at each field site.

**Figure 1.**
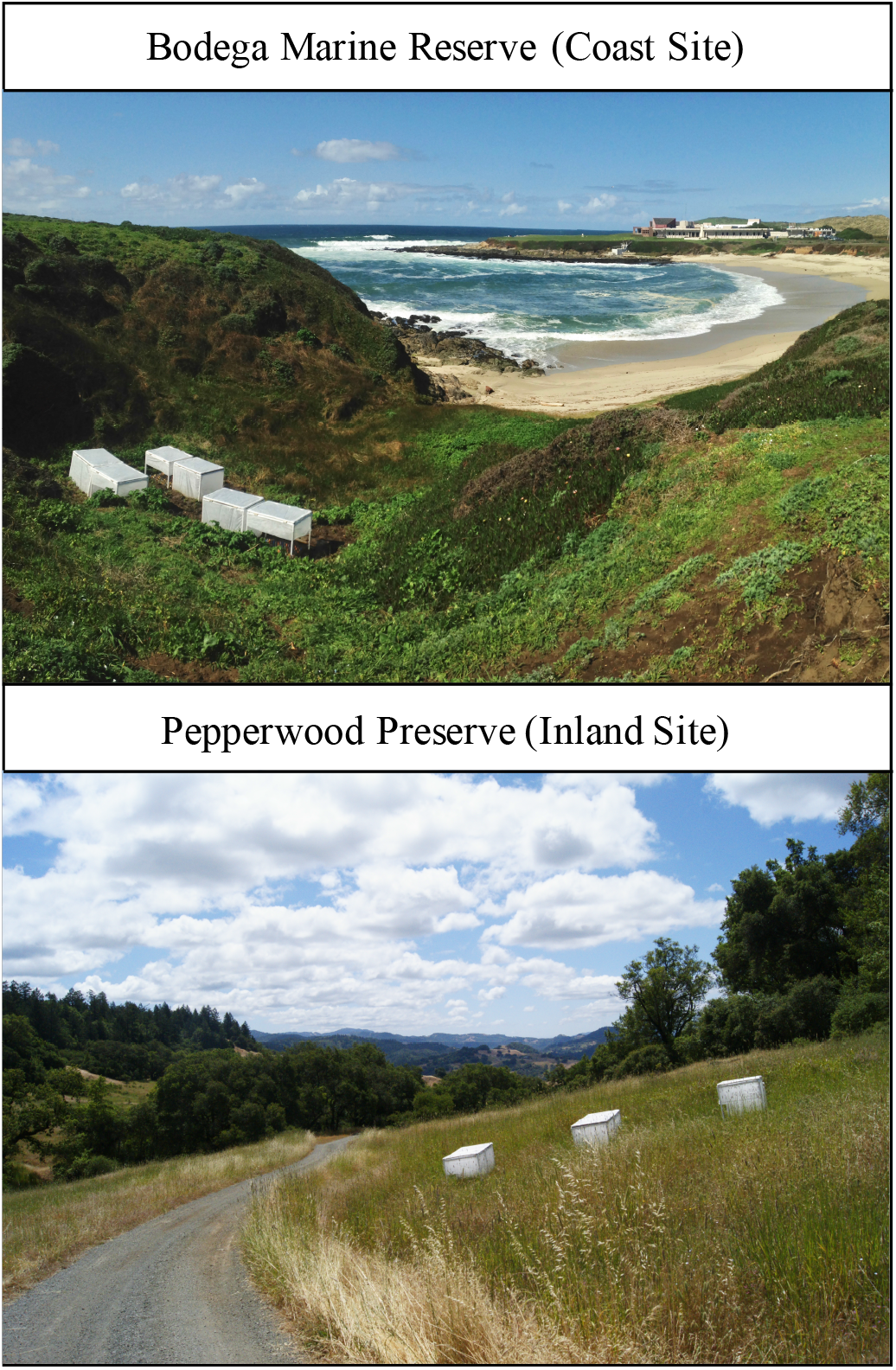
Photos of reciprocal transplant field sites in Sonoma County, CA. The coast garden was located in a perennial seep running from the bluffs overlooking Horseshoe Cove, down to the sandy shoreline of the Pacific Ocean. The inland garden was planted along the margins of a seasonal seep in the Mayacamas Mountains, CA.

All six plots were established with the following dimensions: 216 cm L x 84 cm W (Figure 2B). Plots were positioned haphazardly no farther than 2 meters apart, leaving sufficient room for data collection and plot upkeep. Each plot was cleared of native vegetation prior to transplantation, simulating an artificial landslide event, which are common in California’s headland bluffs (Collins & Sitar 2008). All plots were subdivided into two 108 cm L x 84 cm W subplots (6 per site, 12 total) and randomly assigned a treatment: exclosure or shade control (see details of treatments below).

**Figure 2.**
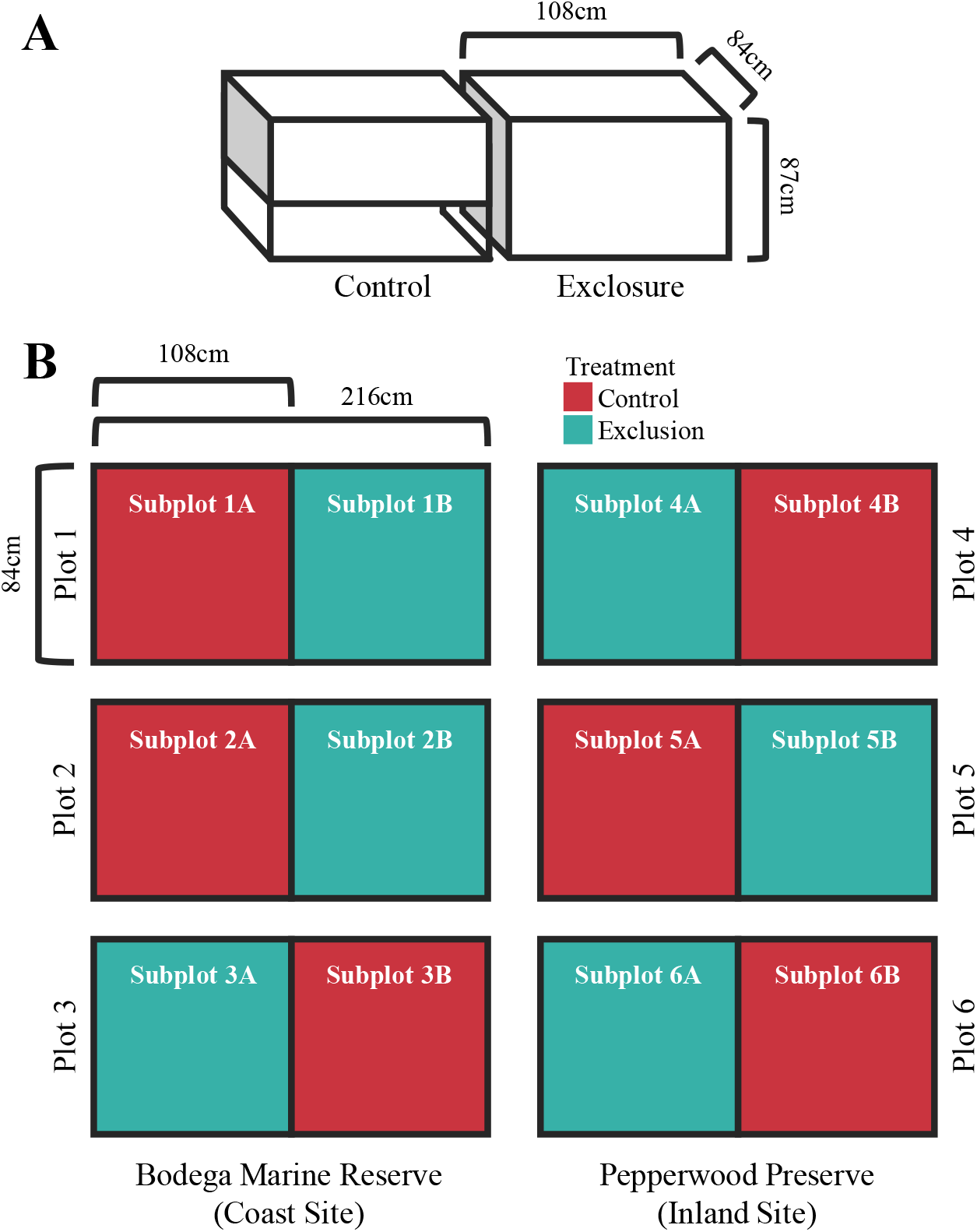
(A) Dimensions of exclosures and shade control shelters used in this study. (B) A diagram of the split-plot design for both coastal and inland sites.

We transplanted biological replicates at both coastal and inland sites a week after the plants were relocated to the BMR greenhouses (March 8^th^ 2017). All *M. guttatus* seedlings utilized in this study were left to mature to the four-leaf stage prior to planting to best ensure transplant success, but still assess field survival prior to reproduction. 25 replicates per population were planted haphazardly in a grid within each designated subplot (100 per subplot, 200 per plot, 600 per site, and 1200 total). In all, 26 individuals died within a week of the initial planting (primarily inland replicates at the inland site) and were replaced immediately. All losses within the first week of planting were considered a result of transplant shock, thus justifying swift replacement.

### Environmental manipulations

To test the combinatorial effects of coastal conditions on population mean fitness, we constructed multiple exclosures with the intent of excluding the aboveground stressors unique to each site – including transient herbivores and oceanic salt spray. Three replicate exclosures (108 cm L x 84 cm W x 87 cm H; Figure 2A) were installed at each site (six total) after planting. PVC pipes were used to construct the scaffold of each shelter, consisting of both a ground level and waist height rectangular quadrat attached at each corner by four PVC legs. All joints were reinforced with PVC cement to improve overall rigidity to withstand wind. A third quadrat was mounted with door hinges on the topside of the scaffold to be used as a lid for plot upkeep and data collection. To ensure that biological replicates were sufficiently buffered from the elements, the lid and scaffold of all exclosures were enclosed using medium weight agrofabric – ordered from OBC Northwest, Inc. (Pro-34 1.0 oz./sq. yd. with 70% light transmission). Agrofabric was also applied to the top two thirds of the shade control shelters, leaving the plants exposed to aboveground stressors, while reducing light in a similar way as the exclosures. All holes made in the agrofabric as a result of the zip ties used to fasten them were reinforced with clear repair tape. Lids were sealed to prevent aboveground herbivore intrusion by installing industrial strength VELCRO brand tape strips along the lip of each exclosure. We buried the exclosures in the ground from 8 - 13 cm, depending on location, to limit herbivore entry through the soil.

### Effects of field manipulations on fitness

Due to concerns by staff of the field reserves about the potential for introgression of nonnative genes into the local gene pools, all plants were regularly emasculated. This practice eliminated the possibility of using flower number or seed set as measures of fitness in this study. Despite this complication, previous studies on local adaptation in this system demonstrates that selection is strong enough for survival to be a sufficient fitness measurement in coastal habitats (Lowry et al. 2008; Lowry & Willis 2010). Thus, to study the effects of habitat and exclosure on the performance of both ecotypes, we collected survival data at seven time points throughout the growing season. All aboveground *M. guttatus* vegetation (dead or alive) of each plant was harvested into brown paper bags at the end of the experiment (June 13^th^ – 15^th^, 2017). These samples were shipped to Michigan State University where they were dried in an oven at 60°C for 2 weeks. Each sample was weighed with an analytical balance to quantify dry aboveground biomass, a valuable but imperfect indicator of plant performance and fecundity (Younginger *et al*. 2017).

To test for differential exposure of aerosolized salt spray across treatments at the coast site, salt traps were designed and installed among all plots roughly following Yura (1997) and Yura & Ogura (2006). In total, 6 salt traps were deployed at the coast site (one per subplot) and 2 traps at the inland site (one per treatment). Traps consisted of a four-sided rectangular plastic prism attached to a cylindrical post. Clear vinyl badge holders were clipped upon each face (one per cardinal direction) to act as protective sheaths for Whatman Brand #2 Qualitative Medium filter paper inserts (cut to size: 8.6 cm L x 5.8 cm W). Badge holders have a native cutout to allow the filter paper to absorb all incident salt spray. Traps were left to collect salt for two weeks starting on May 7^th^, 2017 (24 coastal / 8 inland inserts) and then the filter papers were collected into individual ziplock bags, labelled, and transported back to Michigan State University. Inserts were left to air-dry for 24 hours, placed in Erlenmeyer flasks (1 per trap, 8 total) to soak in 50 ml ultrapure water, and shaken for 1 hour to extract salts. Samples were filtered of all resulting fibers and debris with Whatman No. 44 filter paper and subsequently analyzed for sodium using inductively coupled plasma optical emission spectrometry (ICP-OES; Olesik 1991).

To quantify the degree to which our exclosures affected herbivore activity, we kept a detailed record of how many replicates had experienced any herbivore damage throughout the extent of our experiment. At the end of the season, all plants were categorized as having experienced herbivory or being unscathed by herbivores. Any replicate that died prior to the accumulation of any obvious herbivore related injuries were not counted as having experienced herbivory. We ran a general linear model (GLM) using a binomial distribution to analyze herbivory in the context of presence / absence. Separate models were run for each site. No random effects were evaluated here, as the goal was simply to determine whether the exclosure treatment had indeed effectively reduced the incidence of herbivory at either site.

### Analysis of transplant data

To confirm whether some combination of aboveground stressors contribute to fitness across transplant sites, our data were analyzed using an ASTER modeling approach (Geyer *et al*. 2007; Shaw *et al*. 2008). ASTER is a module developed for the statistical program R that provides a powerful tool for combining multiple fitness components with different probability distributions into a single analysis. The power of ASTER lies in its ability to calculate an expected fitness value for all biological replicates given the order and interdependence of each fitness component. We used ASTER to analyze a composite of 8 fitness components: survival to weeks 1 – 7, all modeled as Bernoulli (0 or 1), and the final harvested aboveground biomass, here modeled as a normal distribution. Due to coding constraints in ASTER, any replicates with a non-zero mass that died before the final observation date (measured post mortem) were scored as a zero for biomass. Likelihood ratio tests were constructed by comparing nested null models to test alternative hypotheses.

We employed a generalized linear mixed modeling approach (GLMM) separately, as an alternative to ASTER because the ASTER module does not readily allow for mixed models with random effects. Thus, the GLMM approach aided us in confirming our ASTER results. Site specific models were developed in lieu of a more comprehensive model, as we were less interested in the effects of site on fitness, but rather the effects of treatment within each site. Response variables included survival to harvest (week 7), modeled using a binomial distribution, and total biomass accrued. Since there is no proper modeling distribution for zero-inflated continuous data in the GLMM framework (data ASTER is quite useful at dealing with), we treated all zero biomass values as missing data and removed them from the analysis. Analysis of the remaining individuals was modeled using a gamma distribution. The following GLMM scaffold was developed for all combinations of site and response (4 models in total): Response ~ Treatment + Ecotype + Treatment:Ecotype + (1|Plot) + (1|Accession). While both ASTER and GLMM modeling approaches have limitations, they complement each other’s weaknesses and together can increase the confidence in interpretation of results.

## RESULTS

### General patterns across field sites and treatments

Overall, 89% of transplants survived to harvest at Pepperwood (inland site) and 73% survived at Bodega Bay (coastal site). Exclosures and control subplots at the inland site saw comparable survival rates, with 90% in the former and 88% in the latter. However, survival diverged markedly at the coastal site with 97% survival in exclosures and 49% in shade control subplots. This vast disparity is driven mostly by discrepancies in ecotype performance dependent on subplot treatment. Plants at the inland site were generally small, having an average biomass of about 0.07g. In contrast, plants at the coastal site had robust growth, ending the season with an average biomass of 1.16g.

### Coastal field site

There were striking differences in the responses of coastal and inland accessions to the exclosure treatment at the coastal field site. At the coast, 87% fewer inland control replicates survived until the end of the experiment than the coastal control plants. The few inland individuals that did survive in the control plants were generally small (Fig. 3A,B). In contrast, the inland plants within exclosures experienced nearly the same survival rates and had a similar mean biomass as coastal replicates under either treatment (Fig. 3A,B). Our GLMM approach found a significance of ecotype x treatment interaction at the coast field site (*P* < 0.0001; Table 2), confirming that differential fitness performance between ecotypes was dependent on treatment.

**Figure 3.**
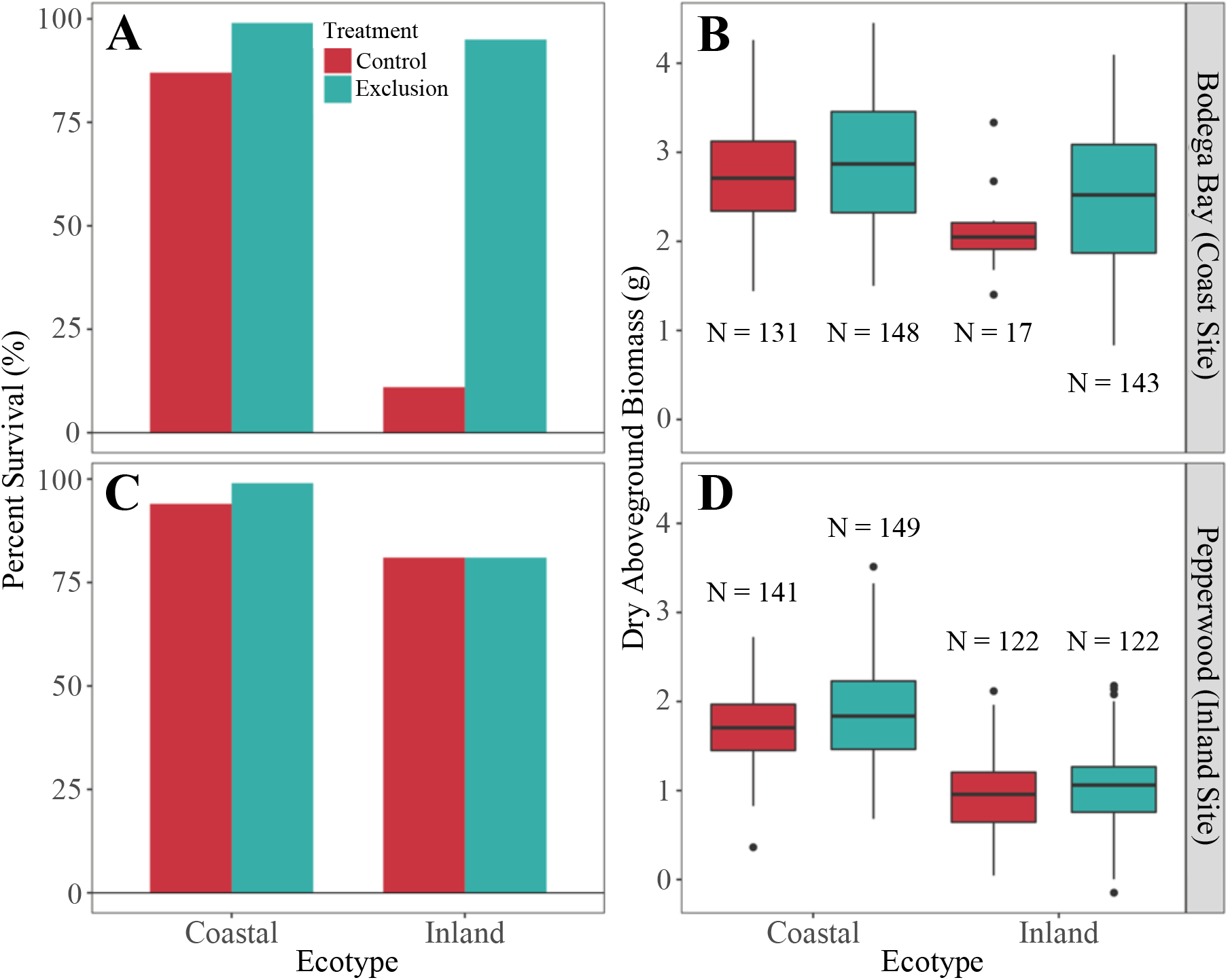
Survival and dry aboveground biomass were measured as fitness proxies in all replicates across both reciprocal transplant gardens to model the effect of ecotype, treatment, and ecotype x treatment interactions on in situ performance. Percent survival is displayed in bar plots specific to the coast (A) and inland (C) sites. Box and whisker plots of dry aboveground biomass at the coast (B) and inland (D) sites for plants that survived to harvest (*N* = sample size of surviving individuals).

**Table 2.** Results for the following Generalized Linear Mixed Model (GLMM): Response ~ Treatment + Ecotype + Treatment:Ecotype + (1|Plot) + (1|Accession). The *P* values for main effects and interactions are provided.

| Site | Response Variable | Treatment | Ecotype | Ecotype x Treatment |
| --- | --- | --- | --- | --- |
| coast | survival to harvest | 0.0005*** | <0.0001*** | <0.0001*** |
|  | dry aboveground biomass | <0.0001*** | 0.10312 | 0.0398* |
| inland | survival to harvest | 0.0337* | 0.0015** | 0.0408* |
|  | dry aboveground biomass | <0.0001*** | <0.0001*** | 0.0308* |

\* $P < 0.05$ , \*\* $P < 0.01$ , \*\*\* $P < 0.001$

ASTER modeling generally confirmed the patterns found by GLMM analysis. Not only were there strong treatment effects at the coastal field sire (*P* < 0.0001; Table 3), but treatment was also found to significantly affect expected biomass for each ecotype (*P* < 0.0001; Table 3). Although the exclosure treatment led to an increase in the expected mean vegetative biomass of both ecotypes at the coastal site, the response of inland plants to the treatment was much more dramatic. This is most evident in Figure 4A, where inland exclosure replicates have accrued an expected mean biomass of 1.58g while inland plants in the control plots had a negative expected biomass (−1.33g) due primarily to low survival.

**Table 3.** Analysis of treatment and treatment-by-ecotype interactions using an ASTER-based modeling approach. Our ASTER models analyzed a composite of eight fitness components, including survival from weeks 1 – 7 and a post-harvest measure of dry aboveground biomass. These components were aligned in the following directional graph in order of general causality: survival to week 1 → survival to week 2 → survival to week 3 → survival to week 4 → survival to week 5 → survival to week 6 → survival to week 7 → biomass accrued. All factors were tested by likelihood ratio tests using nested null models.

|  |  | Null | Alternative | Null | Alternative | Test | Test | Test p |
| --- | --- | --- | --- | --- | --- | --- | --- | --- |
| Site | Factor Tested | df | df | Deviance | Deviance | df | Deviance | Value |
| coast | Treatment | 9 | 10 | -1048.00 | -930.70 | 1 | 117.330 | <0.0001*** |
|  | Ecotype x Treatment | 10 | 11 | -930.70 | -903.31 | 1 | 27.385 | <0.0001*** |
| inland | Treatment | 9 | 10 | -541.45 | -539.24 | 1 | 2.205 | 0.1376 |
|  | Ecotype x Treatment | 10 | 11 | -539.24 | -503.36 | 1 | 35.887 | <0.0001*** |

\*\*\* $P < 0.001$ .

**Figure 4.**
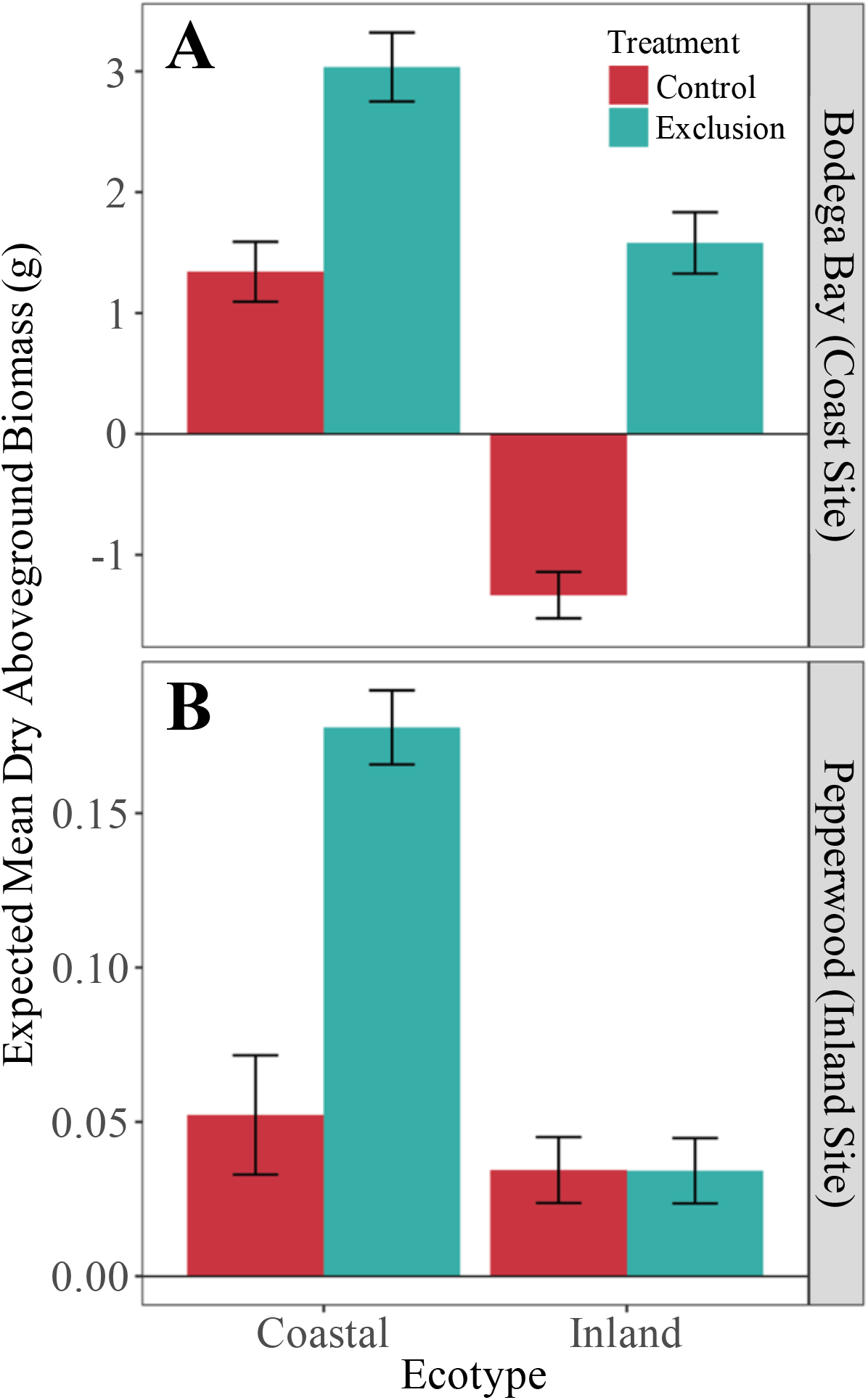
Expected mean dry aboveground biomass of all harvested replicates at the coast (A) and inland (B) sites from the ASTER analysis. Here, expected biomass accounts for the combination of survival with final dry aboveground biomass. A negative expected mean can arise if enough mortality is observed in any particular treatment group. All error bars denote one standard error. Note that scales are different.

### Inland field site

Ecotype, treatment, and ecotype x treatment interaction all had a significant effect on survival and biomass at the inland site (Table 2). However, the magnitude of these differences was relatively small, with a 19% difference in survival between ecotypes and a 3% difference in survival between the treatments. There were significant effects of ecotype on survival (*P* = 0.0015; Table 3) and biomass (*P* < 0.0001; Table 2). Coastal replicates both survived in greater numbers and produced more vegetative biomass than their inland counterparts (Fig. 3C,D). Regardless of these differences, the vast majority plants at this site still survived to harvest.

There was no significant effect of treatment in our ASTER models (*P* = 0.1376; Table 4). However, the ecotype x treatment interaction was significant for biomass (*P* < 0.0001; Table 4). This interaction is the result of an almost three-fold difference in expected biomass between treatments, with exclosure biomass being greater than that of our control plots (Figure 4B).

**Table 4.** Percentage of plants that experienced herbivory across sites and treatments. % Herbivorized was calculated by dividing the final number of herbivorized plants by the total present in each treatment group.

| Site | Ecotype | Treatment | % Herbivorized |
| --- | --- | --- | --- |
| coast | coastal | exclosure | 0 |
|  |  | control | 28 |
|  | inland | exclosure | 0 |
|  |  | control | 29 |
| inland | coastal | exclosure | 0 |
|  |  | control | 7 |
|  | inland | exclosure | 1 |
|  |  | control | 1 |

### Quantifying salt spray and herbivory

Quantifying the concentration of accumulated sodium allowed us to establish whether our exclosures had any significant effect on incident salt spray levels in control vs. exclosure plots. Sodium concentrations were found to be elevated in the ambient conditions of control subplots at the coastal site in comparison to any other combination of treatment and site (Figure 5). Salt samples collected from coastal control subplots had nearly a two-fold higher level of sodium over those sheltered within exclosures. There did not appear to be any difference between the control or exclosure treatments at the inland site. Analysis of a 2% nitric acid blank and clean filter control revealed that our solvent and filter slips had small-to-negligible effects on the sodium content of our test samples. This verifies a pattern of ambient salt reduction as a consequence of our exclosure installations.

**Figure 5.**
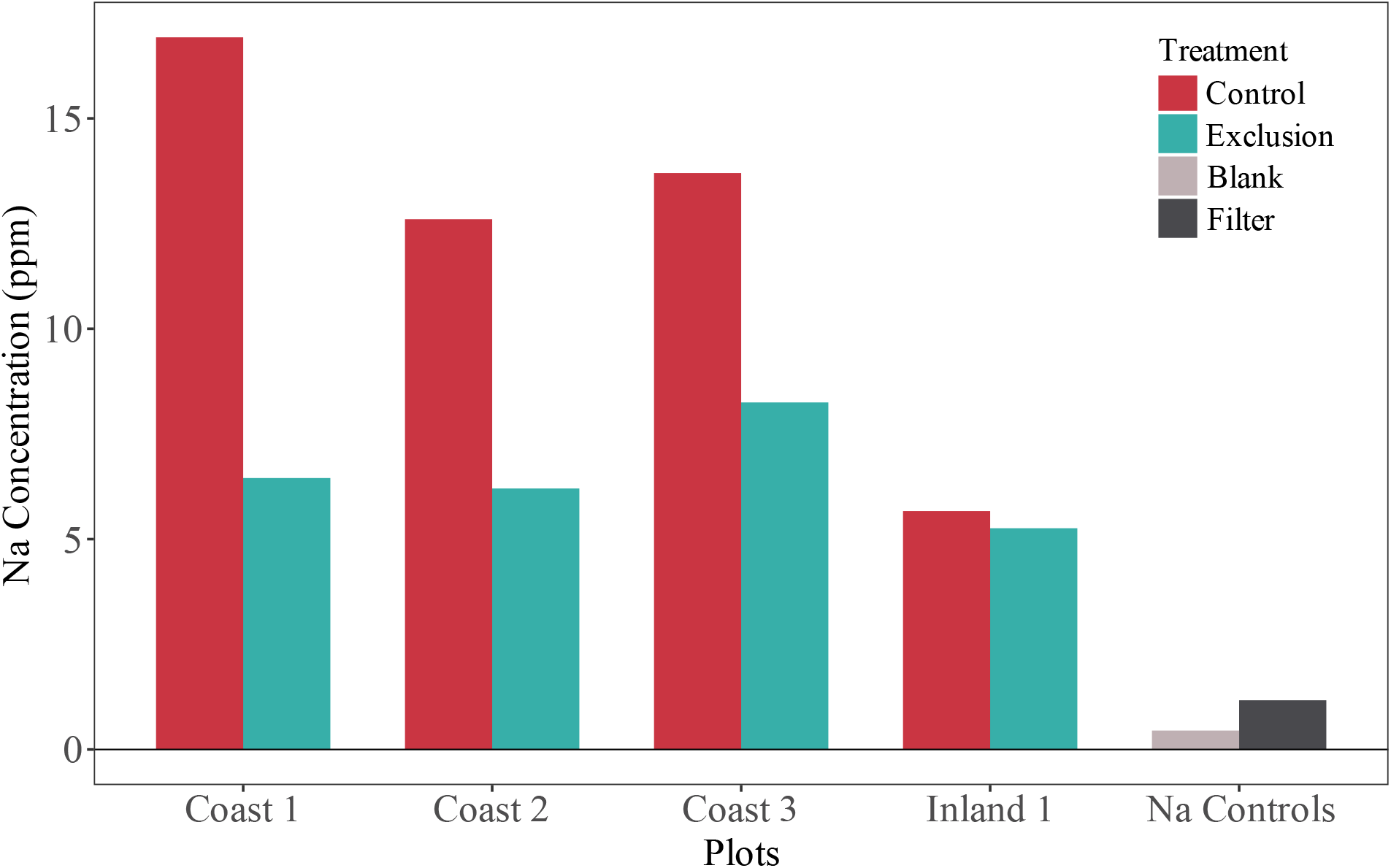
Sodium (Na) concentrations in ppm of all salt trap filter slips left in the field for two weeks, measured using inductively coupled plasma optical emission spectrometry (ICP-OES). Control samples were subject to elements native to each site and plot microhabitat. Exclusion samples were installed within the bounds of their respective exclosure. A blank sample was run consisting only of 2% nitric acid. To gauge whether Na levels were altered in the presence of filter slips, an unadulterated filter control was also analyzed. All coastal plots and one of three total inland plots are shown.

Similar to our sodium analyses, quantifying the incidence of herbivory confirmed the effectiveness of our exclosures. Treatment had a significant effect on herbivory at the coastal site (*P* < 0.0001), nearly eliminating the incidence of herbivore damage to the plants within the exclosures (Table 4). Neither the ecotype nor ecotype x treatment interaction had a significant effect on herbivory at the coast. Though nearly a quarter of coastal and inland controls experienced some herbivore damage at the coast site, there were few observed instances of herbivory at the inland site regardless of treatment (Table 4). In contrast to the coast site, there was no evident effect of treatment on the incidence of herbivory at the inland site (*P* = 0.8105). Only ecotype had a significant effect on rate of herbivore damage (*P* = 0.0189), although how meaningful this was given the low level of herbivory is doubtful (Table 4).

## DISCUSSION

Our results confirm that some combination of aboveground stressors contributes to the adaptive divergence of coastal perennial and inland annual ecotypes of the Seep Monkeyflower, *M. guttatus*. While coastal conditions strongly favor the persistence of native genotypes, nonnative inland accessions at the coastal site were shown to be rescued by exclosure treatments. The effect was so strong that fitness was not merely restored, they thrived at levels comparable to their coastal adapted congeners. Our results suggest that coastal populations have likely adapted primarily to withstand the adverse effects of aboveground selective agents including oceanic salt spray and higher levels of herbivory. In contrast, previous studies have found that local adaptation inland annual populations of *M. guttatus* are likely primarily driven by escape, through earlier flowering, from low belowground soil water availability that occurs during the seasonal summer drought (Lowry *et al*. 2008; Hall et al. 2010; Lowry & Willis 2010). Together, these results demonstrate that different selective agents are responsible for local adaptation at opposite ends of an environmental gradient.

### Patterns of local adaptation

The most significant result of our study was the striking effect of the exclosure on survival and biomass of inland annual transplants at the coastal field site, despite sharing the edaphic conditions of neighboring control replicates. These exclosures ameliorated all of the detrimental effects of natural selection on non-native transplants at the coastal field site. In contrast, the exclosures did not have nearly as great an effect at the inland field site, where coastal perennial plants survived at similar levels as inland annuals. Our analyses demonstrate that a simple control of aboveground stressors in the field can overcome the environmental variables that limit inland fitness at the coastal field site. Thus, some suite of aboveground agents causes the selection that maintains local adaptation of coastal populations of *M. guttatus*.

The effect of our treatment at the inland field site was unexpected, since hypothesized agents of selection like salt spray are not a factor in this habitat and rates of herbivory are much lower. The elevated biomass of the coastal perennial transplants within exclosure were most exaggerated in two of the three inland plots, coinciding with exclosure subplots planted closer to the interior of the seep where soil moisture levels remains favorable for longer. While this result could be an artifact of experimental setup, the same trend was not evident among inland plants. Therefore, it may be that an unknown aboveground set of selective agent limits the performance of coastal transplants in inland habitats. Future studies conducting detailed quantification of herbivory and other factors will be needed to draw any conclusions about mechanism.

In contrast to previous findings on this system, survival of both native and non-native genotypes at the inland site remained comparable throughout the entire growing season. While this result appears to be at odds with local adaptation, it was expected. We have shown in previous studies that despite the consistent performance of coastal transplants early in the growing season at inland field sites, nearly all are killed by the low soil water availability of the summer drought before they have the opportunity to flower (Hall et al. 2006; Lowry et al. 2008; Hall et al. 2010; Lowry & Willis 2010). Fast growing inland annuals survive to flower at very high rates at inland field sites. Thus, by ending the experiment before the summer drought, our results did not capture selection imposed by seasonal drought.

### Salt spray and herbivory as agents of selection at the coastal field site

Salt stress, whether derived from topical incidence or root uptake, can have a range of adverse effects on plants (Boyce 1954; Humphreys 1982; Griffiths 2006). However, coastal populations of *M. guttatus* – often occurring within a few meters of the wavebreak – are known to have a higher tolerance to topical salt application than inland plants (Lowry *et al*. 2008, 2009). The majority of inland replicates in the control plots at the coast site experienced complete vegetative loss consistent with leaf necrosis due to salt spray. Few inland controls persisted long enough to successfully form floral buds. Further, among all bolting survivors, floral stalks and lateral branches turned brown and produced no healthy flowers, likely as a result of salt spray exposure. Similar instances of premature vegetative senescence were noted in inland replicates within coastal exclosures, but this damage was only limited to those tissues in direct contact with the agrofabric. This tissue was presumably experiencing salt stress as a consequence of contact with oceanic salt that had accumulatd on the walls of the exclosure barriers.

Our field observations revealed significantly higher rates of herbivory among control replicates of all genotypes at the coast field site, demonstrating the exclusionary power of our agrofabric treatments. Yet there appeared to be no apparent bias towards one ecotype over another. At least for the field season in which we conducted the experiment, mammalian predation by resident California Voles (*Microtus californicus*) made up a significant portion of damage done to control replicates at the coast – most often evidenced by the complete removal of floral stalks and branches. Though plants in the coastal control plots experienced more herbivory than the plants in the exlosures, it did not appear widespread enough to be the sole explanation for such striking fitness differentials across treatments. Overall, some combination of herbivory and salt spray have clearly contributed to local adaptation of coastal populations of *M. guttatus*. However, we cannot parse the relative importance of these two factors without further manipulative field experiments.

## ACKNOWLEDGEMENTS

The authors thank Benjamin Blackman, Erin Patterson, and the University of California Berkeley greenhouse staff for tending to our germinants, the Pepperwood Foundation Staff, most especially Michelle Halbur, Michael Gillogly, Sonja Barringer, Sloane Shinn, and President/CEO Lisa Micheli for graciously hosting our inland transplants, the staff and faculty of the University of California Davis Bodega Marine Laboratory & Reserve, notably Jacqueline Sones, Kitty Brown, Lisa Valentine, Suzanne Olyarnik, and Director Gary Cherr for providing accommodations for both our coastal transplants and field researchers, to Ruth Shaw, Marcus Warwell, and Christopher Warneke for their attentive modeling advice, and a heartfelt thank you to Kathy Severin for her aid and expertise on the ICP-OES. Comments, insight, and suggestions by William Wetzel, Marjorie Weber, Jeffrey Conner, Diane Ebert-May, Billie Gould, and Nate Emery were essential to the planning and execution of this project. We especially wish to thank Daniel Jackson and Patrick Kearns for their assistance in the field and resilient disposition during the most laborious stretches. Funding was provided by the Michigan State University Plant Science Fellowship and the Lowry Lab Startup Fund.

